# Variability in intrinsic promoter strength underlies the temporal hierarchy of the *Caulobacter* SOS response induction

**DOI:** 10.1101/2025.01.26.634973

**Authors:** Aditya Kamat, Asha M. Joseph, Deeksha Rathour, Anjana Badrinarayanan

## Abstract

Bacteria encode for gene regulatory networks crucial for sensing and repairing DNA damage. Upon exposure to genotoxic stress, these transcriptional networks are induced in a temporally structured manner. A case in point is of the highly conserved SOS response thwat is regulated by the LexA repressor. Studies have proposed that affinity of LexA towards promoters of SOS response genes is the primary determinant of its expression dynamics. Here, we describe an additional level of regulation beyond LexA properties that modulates the SOS response gene expression pattern. Using transcriptomic analyses, we reveal a distinct temporal hierarchy in the induction of SOS-regulated genes in *Caulobacter crescentus*. We observe that LexA properties are insufficient in predicting the temporal hierarchy of these genes. Instead, we find that intrinsic promoter strength underlies the order of gene activation, with differential sigma factor association as a key factor in modulating gene expression timing. Our findings highlight a novel regulatory layer in SOS dynamics and underscore the importance of promoter properties in shaping bacterial stress responses.

## Introduction

The SOS response is a highly conserved bacterial stress response that is activated by a wide range of DNA damaging agents. The damage-inducible nature of the SOS gene network emerges from the regulatory inter-play of two proteins, RecA (activator) and LexA (repressor) [1]. In the absence of DNA damage, promoters of genes under SOS regulation are bound and repressed by LexA [2]. Under DNA damage, a RecA nucleoprotein filament formed on single-stranded DNA triggers the rapid auto-cleavage of LexA, resulting in de-repression and induction of SOS response genes [3–5].

The SOS paradigm is best-studied in *E. coli,* which constitutes over 50 genes, many of which are involved in enabling cell survival under genotoxic stress [6,7]. These include diverse mechanisms that repair or reverse DNA damage as well as regulators of cell cycle progression [8]. Precise regulation of the SOS response is essential due to the costs associated with expression of mutagenic error-prone translesion polymerases, toxin-antitoxin systems and prophages that are also induced when this response is activated [9–12]. Despite the possible fitness costs, the response is important for enabling immediate stress tolerance [13–15] and long-term evolutionary adaptations to a variety of growth environments [16–18].

Several studies over the past years have provided seminal insights into two key features of the SOS response and its regulation via LexA: (i). Temporal hierarchy: SOS response genes exhibit hierarchy in the time taken to reach peak promoter activity, with high-fidelity repair mechanisms peaking earlier as compared to low-fidelity (error-prone) mechanisms [6,19]. Based on this, previous studies suggest a ‘just-in-time’ transcription model for the *E. coli* SOS response timing which mitigates the mutagenic cost of the response via a delay in maximal promoter activity of these genes [19,20]. This phenomenon, observed only at high doses of UV damage, has been attributed to the differential binding affinity of LexA towards promoters of the SOS response genes [19]. (ii). Spontaneous activation: The SOS response is inherently leaky, and pulses of SOS gene expression occur throughout the population even in the absence of genotoxic stress. This intrinsic leakiness in the absence of DNA damage can be attributed to variability in spontaneous LexA cleavage across the bacterial population [21], also contributing to heterogeneity in SOS gene expression.

Along with time to peak expression, time to induction of the SOS response genes may also have significant consequences (especially under lower sub-lethal doses of damage), but this property is less explored. While ‘just-in-time’ transcription organization [22–24] is an elegant method for distributing costs, it is unclear if this is applicable in other bacterial systems, and whether LexA properties alone are sufficient to fully capture the complexities of SOS response induction (such as both induction and peak timing). Indeed, some studies have also implicated other factors in shaping the SOS response dynamics. For example, post-induction, SOS response shows an oscillatory/ pulsatile behaviour in single cells that appears to be modulated by the UmuDC complex in *E. coli* [25]. More recently, cell growth and division rates have been shown to contribute the SOS gene expression heterogeneity, resulting in sub-populations with distinct SOS expression levels and associated growth dynamics [26].

Here we study the temporal dynamics of the *Caulobacter crescentus* SOS response under persistent, sub-lethal doses of DNA damage which allows us to estimate gene expression induction times. We find that genes belonging to the *Caulobacter* SOS response display temporal hierarchy in their time of induction. Our computational analyses reveal that LexA properties do not influence this hierarchy. Instead, we implicate intrinsic promoter strength as a determinant of temporal variation observed in the induction time of the SOS response genes. Our work suggests a role for differential sigma factor affinity as one of the factors shaping the promoter properties and hence, tuning the hierarchy of SOS response induction. This raises the possibility of context-dependent modularity in the SOS response timing that emerges from variation in sigma factor abundance. Together, our work uncovers a simple regulatory feature that influences the timing of the *Caulobacter* SOS response to DNA damage.

## Results

### Characterization of the *Caulobacter* SOS response under mitomycin-C damage

Prior to investigating the temporal hierarchy of the *Caulobacter* SOS response, we first sought to comprehensively characterize the transcriptional response of *Caulobacter* cells to DNA damage. Accordingly, we exposed *Caulobacter* cells to sub-lethal dose (0.25 µg/ml) of mitomycin-C (MMC), a potent inducer of DNA damage (intra-strand crosslinks and DNA mono-adducts) (Fig. S1A). At this concentration *recA* is essential, while survival of wild type cells is not perturbed. We collected samples at 0 min (control), and at 20 and 40 min post-MMC exposure for RNA sequencing (Fig. 1A). At 20 min post-MMC treatment, we observed that 30 genes exhibited increased expression (log_2_FC>1) (Table S4). At 40 min of MMC treatment, this number increased to 71.

**Figure 1.**
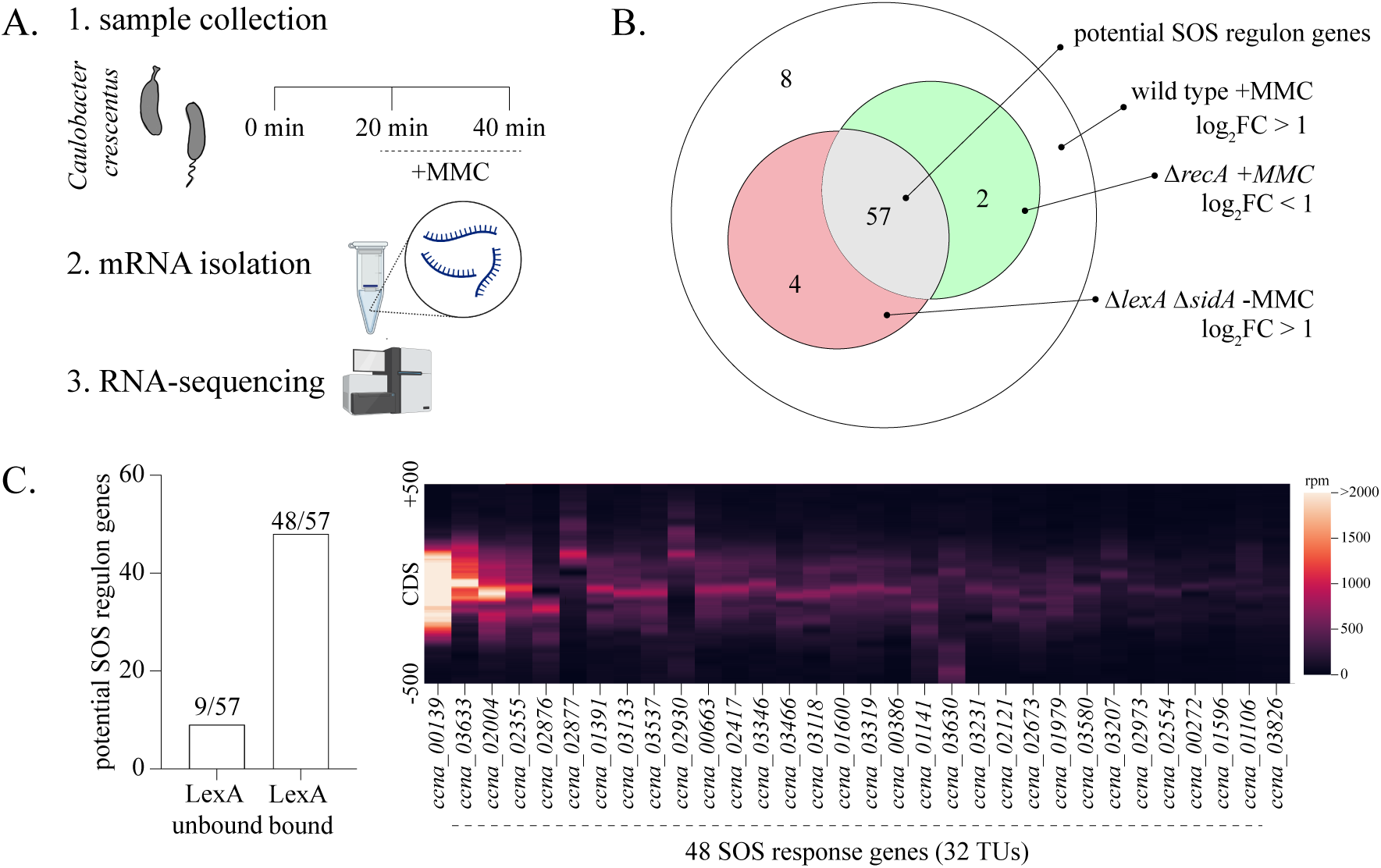
Characterization of the *Caulobacter* SOS response under mitomycin-C damage. **(A)** Schematic summarizing the time course RNA-sequencing experiment. *Caulobacter* cells were treated with 0.25 μg/ml mitomycin-C for 20 and 40 min. Samples were collected for transcriptomic analysis before (0 min, control), and at 20 and 40 min post damage induction. **(B)** Venn diagram representing the genes that meet the listed criteria: 1. Genes induced in wild type cells under MMC damage (white circle). 2. Genes not induced in *ΔrecA* background under MMC damage (green circle) 3. Genes induced in *ΔlexA* in the absence of damage (red circle). Genes fulfilling all three criteria are highlighted in the grey circle. Number of genes in each category is indicated. **(C)** [left] Bar graph indicating whether promoters of the shortlisted genes exhibit binding by the LexA protein as assessed from ChIP-seq analysis [10]. [right] LexA ChIP-seq profile for genes belonging to the SOS response. Normalized reads (in rpm) are represented for 500bp upstream and downstream of the gene CDS. ChIP-seq data were obtained from the GEO database (GSE76721) [10]

To ascertain the proportion of these genes under the SOS response, we set three criteria: 1. Genes regulated by LexA should be induced (de-repressed) in the absence of the LexA repressor even without DNA damage, 2. Gene expression should be dependent on RecA in the presence of damage, and 3. Gene promoter regions should be bound by LexA. For testing the first criteria, we made use of a *ΔlexAΔsidA* background [29,30]. A cell devoid of *lexA* (*ΔlexA*) exhibits extreme filamentation. On the other hand, *ΔlexAΔsidA* cells do not encode the SOS-induced cell division inhibitor (*sidA*) that would be constitutively expressed in cells lacking *lexA.* Hence, this strain background does not yield filamentous cells and significantly rescues the growth phenotype of the *ΔlexA* strain [30]. Out of the 71 genes induced under MMC damage, 61 were found to be under LexA repression, since they were induced in the *ΔlexAΔsidA* background even in the absence of DNA damage (Fig. 1B, Table S4). <u>Genes</u> *sidA* and *lexA* did not exhibit induction in the *ΔlexAΔsidA* background due to poor coverage resulting from their deletion. However, these genes have been previously shown to be under LexA regulation and were thus included in the list of 61 genes de-repressed in the absence of *lexA* [27].

We corroborated these observations by assessing whether these genes were regulated by *recA* as well. We observed that MMC-dependent induction of 57 out of 61 LexA-regulated genes was abolished in a *ΔrecA* background (Fig. 1B). We next tested whether cognate promoters of these potential SOS regulon genes were bound by LexA. For this, we analysed published *Caulobacter* LexA ChIP-seq data [10]. We found that 48 of these genes exhibited LexA enrichment in their promoter sequence. These LexA-regulated genes constituted of 32 transcriptional units (TUs) when we took their operonic arrangements into consideration [27].

Comparison of these genes with a previously annotated *Caulobacter* SOS response (identified via *in silico* analysis [28]) indicated an addition of 20 new genes from our analysis (Fig. 1C). Out of the 35 genes identified to be part of the SOS response in the previous study [28], 7 genes did not qualify our set criteria (Fig. S1B). Six of these genes did not exhibit induction in response to MMC damage (Fig. S1C), lacked LexA ChIP-Seq signal, and displayed no change in gene expression in cells lacking *lexA.* Hence, they are unlikely to be bona fide SOS response genes (Fig. S1C). As an exception, *ccna_02062* (*recN*) exhibited de-repression in the *ΔlexAΔsidA* background but narrowly missed the threshold set for induction under damage (log_2_FC*_recN_*=0.95) (Fig. S1C). We thus exclude this gene from the *Caulobacter* SOS regulon analysis for this study. Collectively, we identify 48 genes (summarized in Table S5) that showcase key features for SOS genes that are directly regulated by RecA-LexA.

### The *Caulobacter* SOS response exhibits temporal hierarchy

Our time course RNA-seq analysis revealed that the *Caulobacter* SOS response is induced in a structured manner. We observed that 18 out of 32 SOS response promoters (∼56%) were induced at 20 min post exposure to MMC (log_2_FC>1) (Fig. 2A). We classified promoters belonging to such genes as ‘early’. The remaining promoters were classified as ‘late’ since they exhibit induction only 40 min post MMC exposure (Fig. 2A). Interestingly, we observed that the promoter regulating the *imuABC* operon associated with SOS-inducible mutagenesis [29] exhibited early induction. On the other hand, *uvrA* gene linked with relatively error-free nucleotide excision repair was induced late upon MMC exposure.

**Figure 2.**
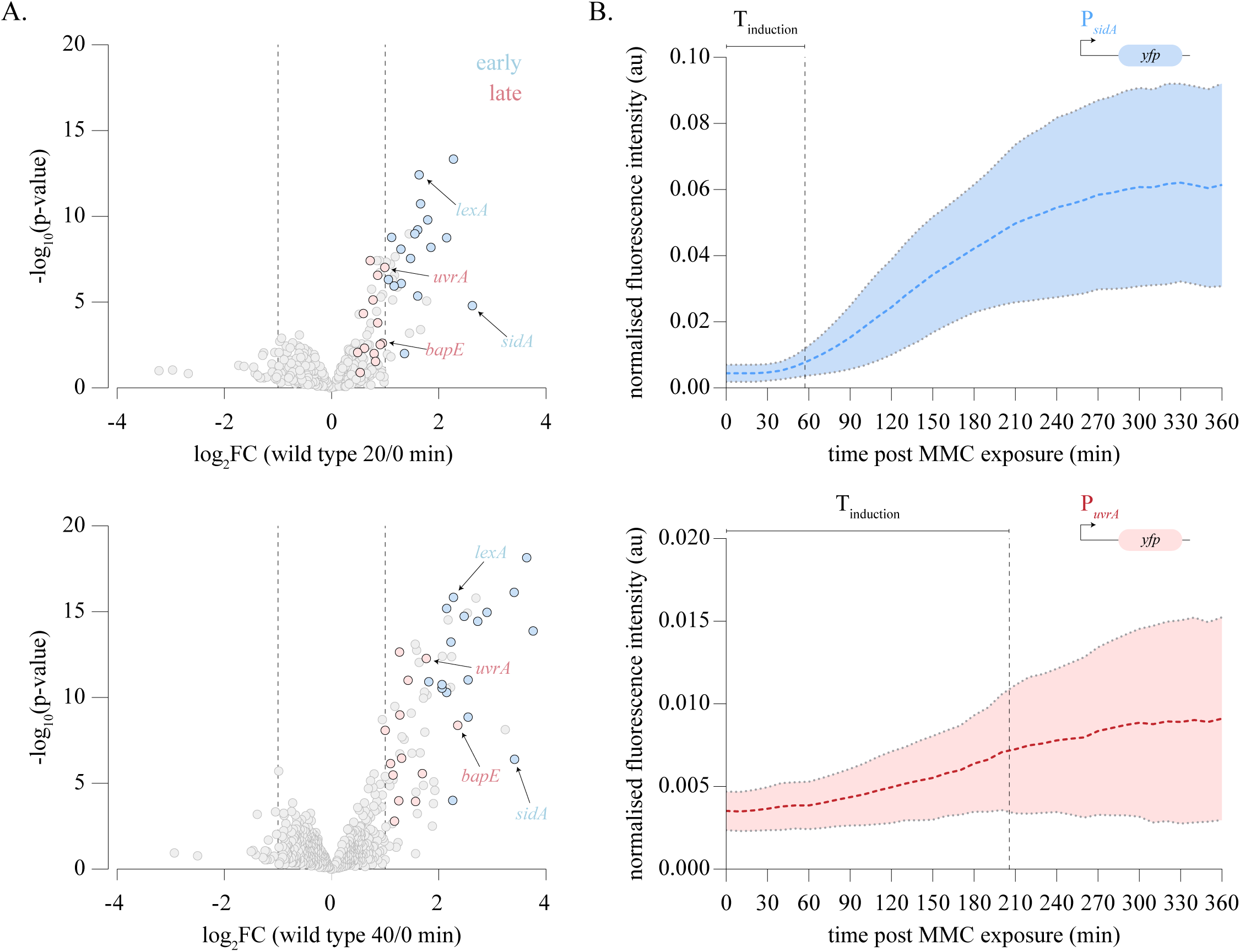
The *Caulobacter* SOS response exhibits temporal hierarchy. **(A)** Volcano plot representing differentially expressed genes for wild type cells exposed to MMC for 20 minutes (top) and 40 minutes (bottom). Early and late SOS genes are highlighted in blue and red respectively. Cut-off for calling a gene as up- or -downregulated is set at log_2_FC>1 and log_10_(p-value)>2 respectively, and are indicated by dashed lines in the volcano plots. **(B)** Fluorescence intensity normalized to cell area for P*_sidA_yfp* [top] and P*_uvrA_yfp* [bottom] cells over 6 hours of MMC exposure. Images were taken every 5 min and intensity over time traces are shown. Dashed line indicates the mean and the shaded region indicates the standard deviation. Time to induction (T_ind_) is indicated on the graph. (0.25 μg/mL, n=25).

To verify these observations, we assessed the induction kinetics of candidate genes for early (*sidA*) and late (*uvrA*) SOS genes using a transcriptional reporter assay. For this, we designed low copy plasmid-based reporters where *yfp* was under the transcriptional control of either the promoter of *sidA* or *uvrA*. Fluorescence intensity normalized to cell area was tracked over time in single cells for these promoter fusion constructs upon exposure to MMC. In line with our RNA-seq analysis, we found that the time to induction of P*_sidA_yfp* preceded that of P*_uvrA_yfp* (Fig. 2B). This phenomenon was independent of damage concentration since the induction of P*_uvrA_yfp* lagged that of P*_sidA_yfp* over a wide range of MMC concentrations (Fig. S2A, S2B). We refer to this phenomenon as the temporal hierarchy of *Caulobacter* SOS response induction, which we note is distinct from timing associated with the peaking of the response.

### LexA properties are not predictive of the temporal hierarchy in SOS response induction

We next investigated the factors that shape this temporal hierarchy of the *Caulobacter* SOS response. Based on studies in *E. coli,* we first assessed whether differences in affinity of the LexA protein towards SOS response promoters could explain the differences in the observed induction times [19]. The degree of divergence of LexA box sequence from the consensus sequence can be scored and has been previously used as a reliable proxy for LexA affinity [30]. LexA boxes with higher divergence exhibit lower affinity as compared to boxes with lower divergence [7,31]. Thus, affinity of LexA towards the SOS promoters can be estimated from variation in the LexA box sequences. Towards this, we first identified cognate LexA boxes for the *Caulobacter* SOS response genes. Using MEME analysis, we identified the LexA boxes for individual SOS response promoters (Fig. S3A, Fig. S3B) [32]. The consensus motif agreed with the previously published motif (GTTCN_7_GTTC) (Fig. S3B) [28].

We conducted principal component analysis (PCA) to test if the LexA box sequences cluster based on temporal gene expression features. The LexA boxes of early and late SOS genes did not show any distinct separation, suggesting no significant difference in their sequences (Fig. 3A). We then scored individual LexA boxes via position weight matrix approach for similarity to the consensus motif. Here too we did not observe any significant difference between the scores for LexA boxes of early versus late SOS genes (Fig. 3B). In further support, we found that the LexA peak heights derived from the ChIP-seq dataset did not vary between early and late SOS promoters (Fig. S3C). Collectively, these results suggest that differential affinity of LexA proteins for early and late SOS promoters may not be the underlying explanation for the observed graded response.

**Figure 3.**
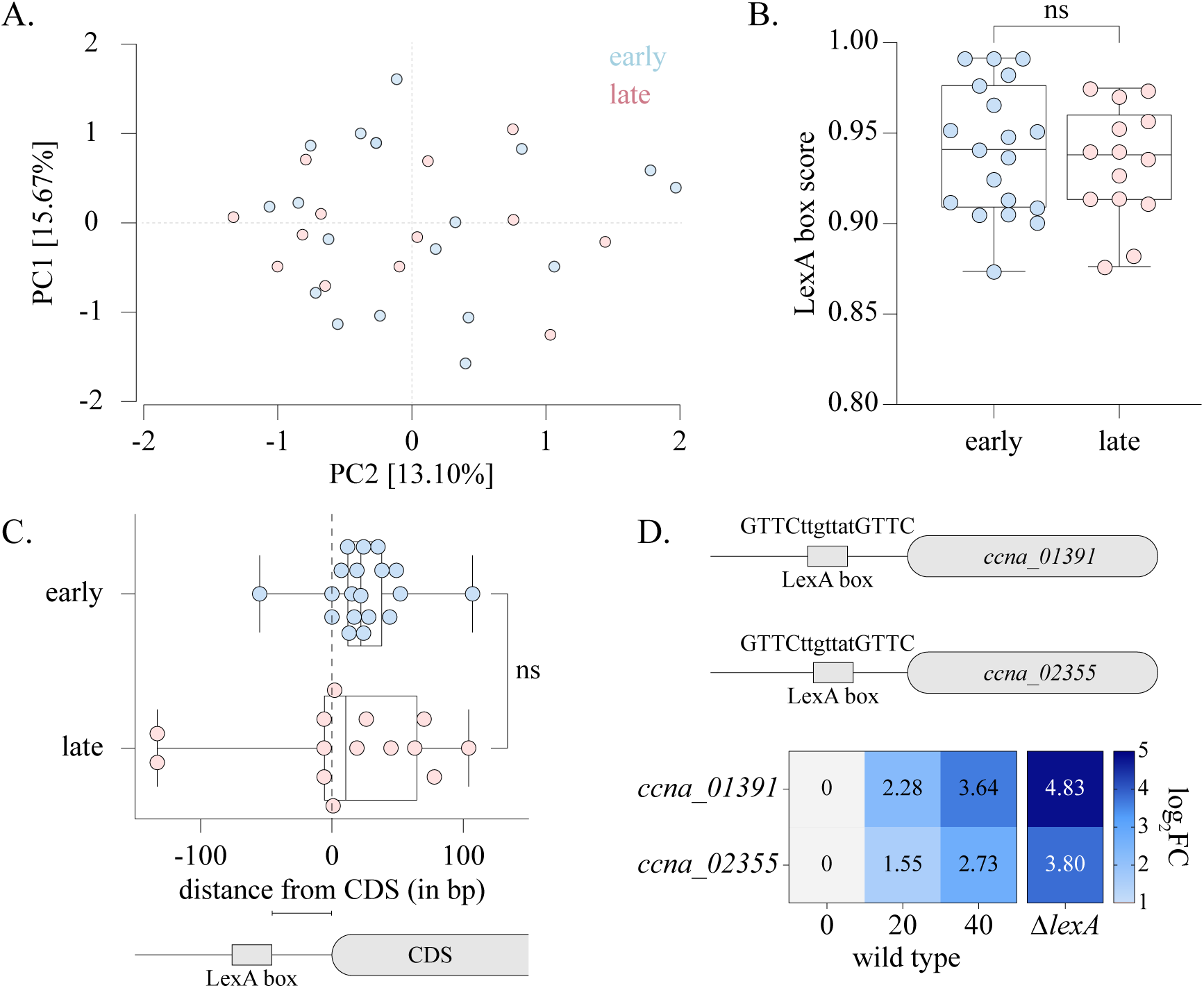
LexA properties are not predictive of the temporal hierarchy in SOS response induction. **(A)** PCA analysis of LexA box sequences for early (blue) and late (red) *Caulobacter* SOS response genes. Percentage variance explained by each compartment are indicated on the axes. **(B)** Box and scatter plot representing scores of LexA box motifs belonging to early and late SOS response genes. Upper and lower limits of the box represent the 25^th^ and 75^th^ percentile of the data, line within the box denotes the median and the box whiskers represent the upper and lower extremes of the data (for this and all other box plots in this study). Scores for individual LexA boxes of the early and late genes are represented by blue and red points respectively. t-test (unpaired), n.s. – not significant. **(C)** [top] Box and scatter plot indicating distance of LexA box motif from the CDS of early (in blue) and late (in red) SOS response genes. Mann Whitney test, n.s. – not significant. [bottom] schematic representing calculation of distances between LexA box motifs and its cognate CDS. **(D)** [top] *ccna_01391* and *ccna_02355* possess identical LexA box motifs. [bottom] Heat map of log_2_FC values from RNA-seq experiments for *ccna_01391* and *ccna_02355* over 0, 20 and 40 minutes post MMC exposure in wild type, and *ΔlexAΔsidA* (no DNA damage) are shown. Fold change value is shown within each square.

We noted that a majority of SOS response promoters (∼97 %) carried only a single LexA box. We wondered whether features of the LexA boxes other than sequence or copy number could contribute to the SOS response timing. We thus assessed the distance of the LexA box from its CDS. We found that the distribution of LexA box distance from gene start was similar for early or late SOS genes (Fig. 3C). Other LexA box features such as spacer length or spacer GC content were invariant as well (Fig. S3A, S3D). Thus, LexA box properties do not appear to be good predictor of SOS response temporal hierarchy.

In support, we identify two SOS response genes, *ccna_01391* and *ccna_02355* which possess identical LexA box motifs (GTTCttgttatGTTC) (Fig. 3D). Despite this, our time course RNA-seq experiment revealed that the induction kinetics of these two genes under MMC exposure were significantly different (Fig. 3D). Moreover, in the absence of *lexA* as well, these genes exhibited variability in their induction levels. Thus, identical LexA sequences can give rise to varied induction kinetics, suggesting the need to look beyond LexA-associated properties of the SOS response genes.

### Intrinsic promoter properties contribute to temporal hierarchy in SOS response gene induction

Our observations so far suggested that variation in LexA box properties did not determine the temporal hierarchy of the *Caulobacter* SOS response. Given that two SOS response genes with identical LexA boxes showed differences in their induction levels even in the absence of *lexA,* we wondered whether variation in intrinsic promoter properties was a determinant of the observed temporal hierarchy. In this direction, we assessed whether transcription unit (TU) length or gene distance from the origin of replication were sufficient in explaining temporal differences of the SOS response genes. We found that neither of these properties were different between early and late-induced genes (Fig. 4A-B). We next tested whether variation in promoter strength shaped the temporal hierarchy of the *Caulobacter* SOS response. Previously, studies have implicated a role for differential affinity of sigma factors towards its cognate promoters as a source of variance in promoter activity [36,37]. We thus asked whether sigma factor association was different between early and late gene promoters. For this, we analyzed published ChIP-seq data of RpoD (the principal sigma factor, σ^73^) with the SOS response promoters [34,35]. We found that RpoD associated with 89% of the early SOS gene promoters (Fig. S4A, Table S6). However, only 47% of late SOS gene promoters had RpoD enrichment in their promoter sequences, with ChIP-seq profiles also showing lower binding affinity when compared to RpoD bound at early gene promoters (Fig. S4B, Table S6). We wondered whether late gene promoters are instead associated with alternate sigma factors. In this direction, we found that two late SOS gene promoters (*ccna_00139* (*chlI*), and *ccna_02673* (*uvrA*)) appeared to be under the regulation of the alternative sigma factor σ^32^ (RpoH). Both genes showed upregulation in an RNA-seq experiment of a *Caulobacter* strain over-expressing a stable mutant of σ^32^ (*rpoH^V65A^*) (Fig. S4D) [36]. Moreover, analyzing σ^32^ ChIP-seq data also revealed σ^32^ enrichment at the *ccna_02673* (*uvrA*) promoter (Fig. S4B) [35]. On the other hand, none of the early SOS promoters showed σ^32^ enrichment or upregulation upon σ^32^ overexpression (Fig. S4C-D, Table S6).

**Figure 4.**
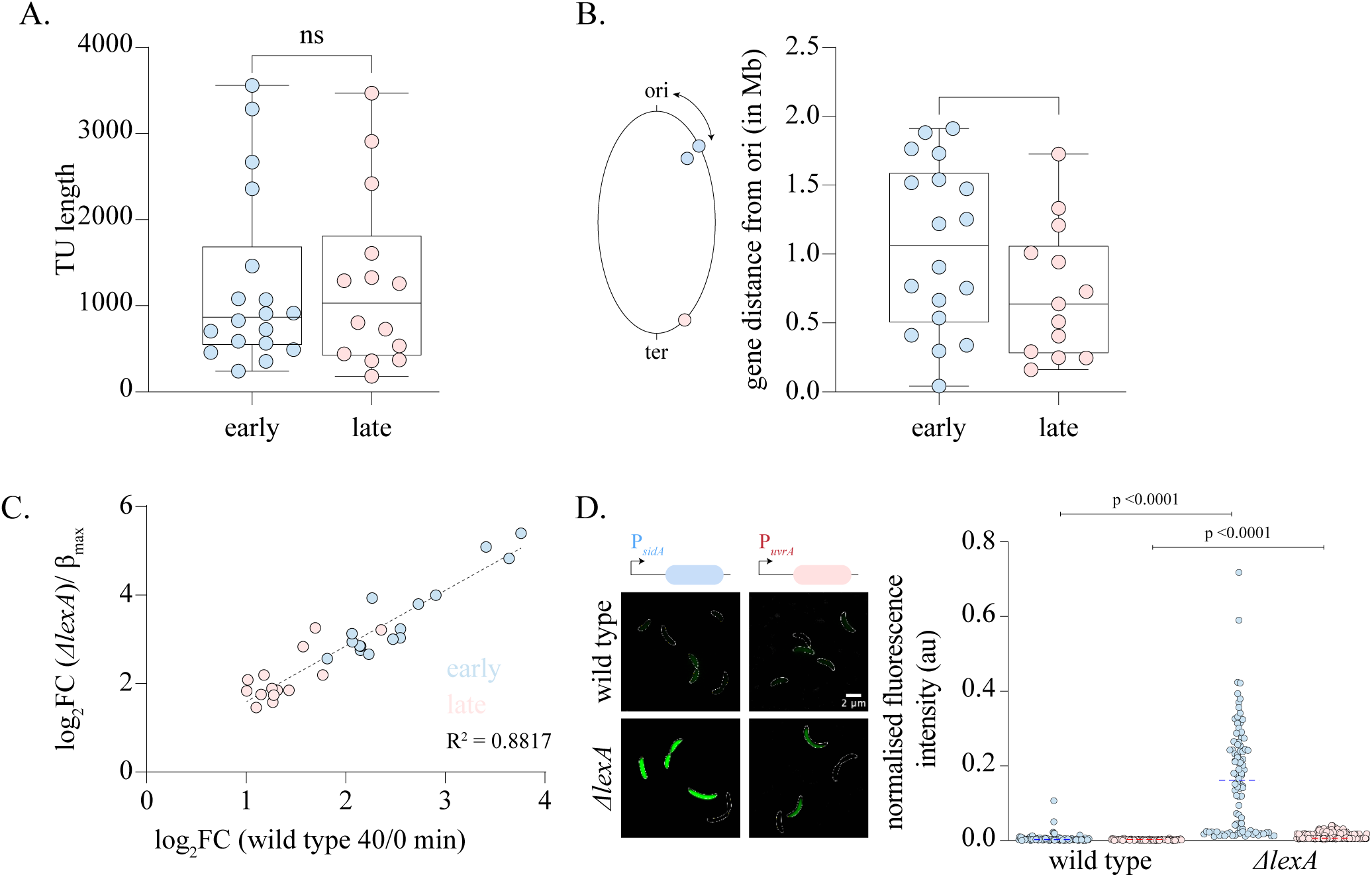
Intrinsic promoter properties contribute to temporal hierarchy in SOS response gene induction. **(A)** Box and scatter plot indicating TU lengths of early (blue) and late (red) SOS response genes. XY test, n.s. – not significant. **(B)** [left] Schematic representing the calculation of gene distance from *Caulobacter* replication origin. [right] Box and scatter plot indicating distance of *ori* from the CDS of early (blue) and late (red) SOS response genes. t-test (unpaired), n.s. – not significant. **(C)** Log_2_FC values for genes in wild type cells upon 40 min of MMC exposure are plotted against Log_2_FC for *ΔlexAΔsidA* cells (representing intrinsic promoter strength/ β_max_) in the absence of DNA damage. Values for early SOS response genes are highlighted in blue while values for late genes are highlighted in red. Pearson correlation r^2^ 0.8817 CI[0.8746 to 0.9708] p-value < 0.0001. **(D)** [left] Representative cells showing induction of P*_sidA_-yfp* (candidate early gene) and P*_uvrA_-yfp* (candidate late gene) in wild type and *ΔlexAΔsidA* backgrounds. [right] Scatter plot show fluorescence intensity distribution normalized to cell area for the respective transcriptional reporters (n=100). Significance was estimated via the Mann Whitney test.

We next assessed promoter strength variation across the entire SOS response to test whether there was a relationship between promoter strength and the observed temporal gene expression hierarchies. Promoter strength of a gene (β_max_) is defined by its maximal expression rate [33]. For SOS response genes, expression rate is dependent on LexA levels and hence expression rate would tend to β_max_ in the absence of LexA protein. Thus, promoter strength for SOS response genes can be estimated based upon expression in the *ΔlexAΔsidA* background. Transcriptome analysis of the *ΔlexAΔsidA* strain revealed that the fold change values of SOS response genes exhibited a strong correlation with their order of induction (hierarchy) (R^2^=0.89), with early induced genes showing higher maximum expression (promoter strength) as compared to genes induced late (Fig. 4C). We corroborated these results with promoter fusion experiments in the *ΔlexAΔsidA* strain and observed a similar trend for our candidate gene promoters for early (*sidA*) and late (*uvrA*) SOS genes (Fig. 4D), with higher *yfp* expression from the *sidA* promoter when compared with the *uvrA* promoter. Along with temporal hierarchy, another hallmark of the *E. coli* SOS response is heterogeneity in gene expression. Strikingly, we also noted that there was significant heterogeneity in the gene expression profiles of *ΔlexAΔsidA* cells. Comparison of the level of variance from mean (coefficient of variance, CV) for P*_sidA_* and P*_uvrA_* promoters in wild type cells revealed a high level of heterogeneity in expression from both promoters in the absence (leaky expression) as well as presence of damage (Table S7; CV 0.76 and 0.53 for P*_sidA_* in the absence and presence of damage respectively. CV 0.55 and 0.76 for P*_uvrA_* in the absence and presence of damage respectively). This heterogeneity was also seen in cells lacking the repressor *lexA*, with CV of 1.3 for P*_sidA_* and CV of 0.9 for P*_uvrA_*. In sum, two key inferences can be drawn from these observations: a. Hierarchy and heterogeneity in gene expression of the *Caulobacter* SOS response is observed even in the absence of the LexA protein. b. The hierarchy is correlated with the promoter strength (Fig. 4C, 4D). Taken together, our data suggest that intrinsic promoter strength, possibly shaped by differences in sigma factor binding, contributes to the temporal hierarchy of *Caulobacter* SOS response induction.

## Discussion

In this study, we investigate the temporal hierarchy observed in the induction of *Caulobacter* SOS response genes. Our work reveals the role of promoter strength as a determinant of this hierarchy. It is possible that such promoter strength variation forms a fundamental regulatory layer for the development of temporal hierarchy in this pathway across organisms.

Collectively, previous studies together with our present findings underscore the presence of hierarchy in two temporal features of the SOS response i.e., time to induction and time to peak response. However, both these features exhibit some contrasting properties: i. Variation in time to peak response has been observed distinctly only at higher concentrations of UV damage [19], while hierarchy in the time to induction can be observed even at sub-lethal concentrations of DNA damage. ii. Error-prone SOS genes seem to peak later as compared to error-free genes [6,19]. On the other hand, at least in the case of *Caulobacter*, error-prone *imuABC* genes are induced earlier than error-free *uvrA*. iii. These temporal features differ in their underlying regulatory logic. While peak time is determined by the affinity of the LexA protein towards the SOS promoters [19], the rate of induction is independent of LexA. Going forward, it will be important to dissect the relevance and contribution of these modes of temporal regulation for survival under DNA damage.

### A role of intrinsic promoter strength and growth context in shaping the SOS response

The regulatory landscape of *E. coli* is estimated to have >4000 genes in turn regulated by >200 regulators [37,38]. How do bacteria determine what genes to turn on and when? A crucial determinant of the bacterial transcription landscape is based on the modularity imparted to the RNA polymerase subunit by the sigma factors [39]. During exponential phase, most genes are under the regulation of σ^70^ (RpoD) [40]. Bacteria also possess alternative sigma factors that are dedicated to activate regulon genes under specialized conditions [39–41]. In our experimental regime, we maintain *Caulobacter* cells in exponential phase. Thus, we hypothesized that the SOS response genes would be under σ^70^-based regulation. Instead, we found that σ^70^ exhibits differential enrichment, predominantly at the early SOS genes as compared to the late SOS genes. Some of the late genes did also exhibit regulation by the alternative sigma factor, σ^32^ (RpoH) [42,43]. Thus, regulation by different classes of sigma factors may contribute to the temporal hierarchy in gene induction observed in the *Caulobacter* SOS response. Interestingly, sigma factors have been associated with the emergence of temporal hierarchy in flagellar synthesis pathway of bacteria as well [44,45].

Indeed, several stressors such as oxidative stress, heat stress, nutrient starvation, stationary phase and antibiotic treatment are also known to trigger the SOS response [46–49]. It is possible that a different hierarchical order might be crucial for survival under such conditions. It is tempting to speculate a context-dependent modularity in the SOS response temporal hierarchy based on the sigma factor involved. In support of this hypothesis, the RpoH-regulated late genes of the SOS response exhibit higher expression levels as compared to the RpoD-regulated early genes in an RpoH^V65A^ over-expression background (Fig. S4C) [36].

### Timing the SOS response

Is temporal hierarchy essential for bacterial survival under DNA damage? As discussed above, variation in peak time has been hypothesized to temporally segregate repair mechanisms based on their mutagenic potential [50]. Thus, error-prone polymerases such as UmuC have been observed to exhibit peak response times much later as compared to genes with no mutagenic cost associated with their induction [6,19]. However, this is unlikely to insulate individual bacterial cells from the cost of mutagenesis. Even though error-prone genes response peak late, they exhibit robust induction early, and at the same time as the error-free genes [6]. For example, the UmuDC operon of *E. coli* is induced 5-fold within 5 min of exposure to UV damage [6]. It is possible that there is mutagenesis associated with the early induction of error-prone polymerases in a sub-population of cells. While there may be costs at the single-cell level, this may serve a crucial function in enabling evolvability and population survival. In contrast, when considering time to gene induction we find that *bapE,* which promotes damage-induced apoptosis, is induced late [51]. Delaying the expression of *bapE* may enable cells to prevent premature commitment to cell death, and prioritize DNA repair and cell survival instead. However, in light of the role of different sigma factors in determining induction times, it is possible that hierarchy may be subject to change depending on the stress and growth conditions. Our work highlights the need to further investigate the temporal hierarchy of the SOS response in the context of different stressors and growth conditions that bacteria may be likely to encounter in their natural environments.

### Limitations of present study and future outlook

In this study we were unable to map the association of many other sigma factors with SOS promoters due to the lack of information on their binding properties. Additionally, we failed to generate transcriptional reporters with LexA binding sites swapped between early and late SOS promoters due to the challenging nature of the sigma factor binding sites and their organization with respect to the LexA binding sites. Such experiments could provide insights to the importance of maintaining hierarchy for SOS function. Finally, recent work has highlighted heterogeneity in the SOS response at the single-cell level. We too observe the same in case of our exemplar ‘early’ and ‘late’ promoters. It is crucial to understand how the population-level structuring of SOS response timing plays out in single-cells (and across multiple gene promoters) to assess the effects of the same on cell survival. The development of single-cell RNA-seq techniques in bacteria, combined with single-cell microscopy will be a powerful method for future investigations in this direction [52,53].

## Materials and methods

### Bacterial strains and growth conditions

All strains, plasmids and oligos used in this study are listed in Table S1-S3 respectively. *Caulobacter crescentus NA1000* cells were cultured in peptone yeast extract (PYE) media at 30°C while shaken at 200 rpm. Antibiotic (kanamycin), DNA damaging agent (MMC) were introduced into the culture at suitable concentrations, wherever required.

### Fluorescence microscopy and analysis

Cultures of *Caulobacter* were grown overnight in PYE media supplemented with kanamycin (5 μg/mL). For time lapse microscopy, the cultures were back-diluted to OD_600_ of 0.025. 3 hours post back-dilution, when OD_600_ reached ∼0.1, 2 μL of culture was spotted on 1.5% low-melting GTG agarose pads supplemented with PYE, kanamycin and required level of MMC damage. Cells were then imaged over 6 hours with an interval of 10 minutes in an OkoLab incubation chamber maintained at 30°C. All imaging was performed on a Nikon Eclipse Ti-2E epifluorescence microscope equipped with a 60x/1.4 NA oil immersion objective, a motorized stage and an LED light source (pE-4000). The *yfp* reporter constructs were imaged with an exposure time of 500ms (λ =490nm, 50% LED power) by an Orca Flash 4.0 camera. Focus was ensured over the course of the time lapse by the Nikon Perfect Focus System. Oufti software was then used for cell segmentation and estimation of normalized fluorescence intensity over time [41]. In order to calculate induction time (T_ind_)for the reporter constructs, we fit a logistic curve model (R^2^ > 0.99) to the mean of each induction profiles in Matlab. Induction time was then estimated as the time required for doubling of the mean initial fluorescence intensity (at time=0). For estimating coefficient of variation (CV), standard deviation of fluorescence intensity normalised to cell area for a given dataset was divided by the mean of that dataset.

### RNA-sequencing

#### Sample collection

Overnight cultures of *Caulobacter* grown in PYE media were back-diluted to 0.025 OD_600_. When OD_600_ reached ∼0.1, 0.25 μg/mL MMC damage was introduced in to these cultures. *Caulobacter* cells were harvested (2mL/sample) for RNA-seq at 0, 20, 40 minutes post damage exposure. The samples were then centrifuged at 10000g for 5 minutes following which supernatant was discarded and the pellets were snap frozen and stored at -80°C until further processed. Data for wild type no damage control was derived from four biological replicates, all other conditions were repeated twice.

#### mRNA isolation and sequencing

mRNA was isolated from the collected pellets as per details mentioned in a previous study [42]. Pre-heated trizol (65°C) was added to each pellet and cells were lysed using a Thermomixer (65°C, 2000rpm, for 10 minutes). Total RNA was extracted from the lysed cells using the Direct-zol RNA MiniPrep kit. Any DNA contamination was then removed via DNAse treatment and RNA was purified using RNA Clean & Concentrator-25 kit. Integrity of the total RNA was tested by the Tapestation instrument. Ribosomal RNA was then depleted using the Thermo RiboMinus kit. mRNA was purified using RNA Clean & Concentrator-5 kit. The mRNA samples were submitted for sequencing to the NCBS next generation sequencing facility. NEBNext Ultra II Directional RNA Library Prep kit was used for library preparation and samples were sequenced via the Illumina NovaSeq 6000 platform.

#### RNA-seq analysis

RNA-seq results were obtained as raw reads in fastq format. The raw reads were mapped to the genome of *Caulobacter crescentus* NA1000 strain (accession number: NC_011916.1) using the Burrows–Wheeler Aligner (BWA) method (raw read quality > = 20). Samtools was used to filter out any multiply mapped reads. Read count per gene was then calculated using Bedtools [54]. Normalization and differential gene expression analysis of these datasets were carried out using the EdgeR package [55]. Genes with log_2_FC values > 1 were denoted as differentially expressed genes. All analysed data are provided in Table S4, and raw data are deposited in GEO. Graphs were plotted in Graphpad prism (10.4.1), and statistical tests were carried out in the same software.

### Survival assay

Cultures of *Caulobacter* grown overnight in PYE were back-diluted to OD_600_ of 0.075. Three hours post back-dilution, all cultures were normalized to OD_600_ 0.3, and diluted serially in 10-fold increments (10^−1^–10^−8^). 6 μl of each dilution were spotted on PYE plates containing no DNA damage (control) or MMC damage (0.25 µg/mL). The plates were then incubated for 48 hours at 30°C post which the plates were imaged, and survival was based upon the number of spots grown. All survival assay experiments were repeated twice.

### Analysis of ChIP-seq data

Raw reads for ChIP-seq data for *Caulobacter* LexA, RpoD and RpoH were obtained in fastq format from previous studies [10,36]. These data were analyzed as described previously and briefly described ahead [56]. The raw reads were aligned to the *Caulobacter* genome (accession number: NC_011916.1) using the Burrows–Wheeler Aligner (BWA) tool. Aligned reads were then converted into the .bed format. Bedtools was used in order to estimate coverage at every nucleotide position. These normalized reads were smoothened by applying a Gaussian filter in Matlab, in order to plot individual ChIP-seq profiles for RpoD and RpoH. Peak calling from the analyzed RpoD data was conducted using the Peakzilla software [57]. RpoD peaks were assigned to SOS response genes if their predicted peak summit position was located within +/- 100 bp of the gene CDS. For calculation of LexA peak heights, maxima of normalized reads counts were estimated.

### Bioinformatic analysis of LexA box properties

#### Identification of LexA box sequences of the SOS response genes

For identifying LexA boxes in the SOS response promoters, DNA sequences 200bp upstream and 75bp downstream of the *Caulobacter* SOS response gene CDS was subject to analysis by the MEME discovery program [32]. The program identified motifs within the provided sequences that repeated one or more times. All identified LexA boxes from the program are summarized in Fig. S3A.

#### Estimation of LexA box scores

In order to score individual LexA box sequences with respect to the consensus motif, a position weight matrix (PWM) was generated using the Biostrings package. The PWM score was then calculated using the “PWMscoreStartingAt” command.

#### Calculation of SOS gene distance from origin

The location of *Caulobacter crescentus NA1000* origin was retrieved from the DoriC database [58,59]. We estimated distance from the origin by calculating the shortest distance of the OriC from the start position of the respective gene.

## Author contributions

AK and AB conceived the project. AK led the project, carried out majority of the experiments, generated tools and conducted data analysis. AJ carried out RNA-seq experiments. DR contributed to strain construction. AB supervised the project and procured funding. AK and AB wrote the manuscript, with feedback from all authors.

## Acknowledgements

Authors are grateful to Prof. Meriem El Karoui and Prof. Andrea Weisse for helpful feedback and advice on the work. The authors acknowledge Neha Sontakke for development of analysis methods for the RNA-seq data. This work was supported by funding to AK (TIFR graduate student fellowship) and to AB via the India Alliance Intermediate Grant (Grant number IA/I/21/1/505630) and intramural funding via NCBS-TIFR (Grant number 03/3/2019/R&D-II/DAE/4749).

## Data availability

All sequencing data are deposited in GEO database.

## Supplementary figure legends

**Figure S1.**
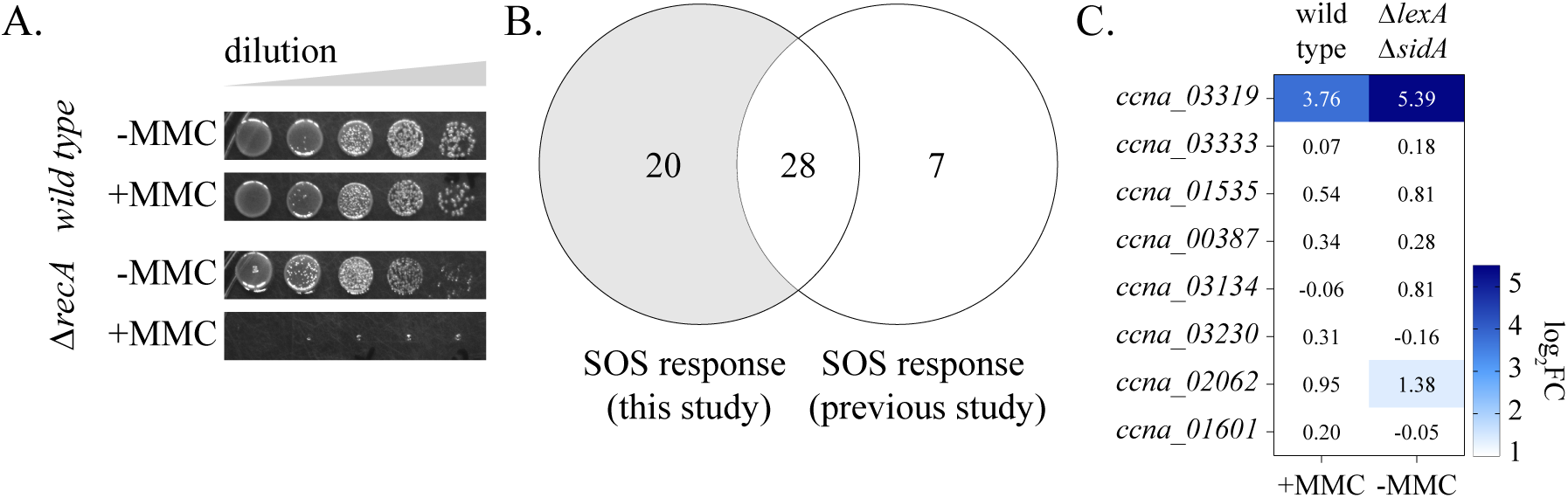
Characterization of the *Caulobacter* SOS response under mitomycin-C damage. **(A)** Survival assay of wild type and *ΔrecA* cells in the presence and absence of MMC (0.25μg/mL). Representative image shown from three biological replicates. **(B)** Venn diagram comparing SOS response genes identified in this study with the *in silico* analysis by da Rocha et al., 2008. Number of genes in each category are indicated. **(C)** Heat map for log_2_FC values of genes previously identified as *Caulobacter* SOS response genes which do not fulfil criteria put forth in this study. Log_2_FC values for an SOS-induced gene (*ccna_03319)* in wild type cells exposed to MMC damage for 40 minutes and for *ΔlexAΔsidA* cells in the absence of damage are shown for comparison.

**Figure S2.**
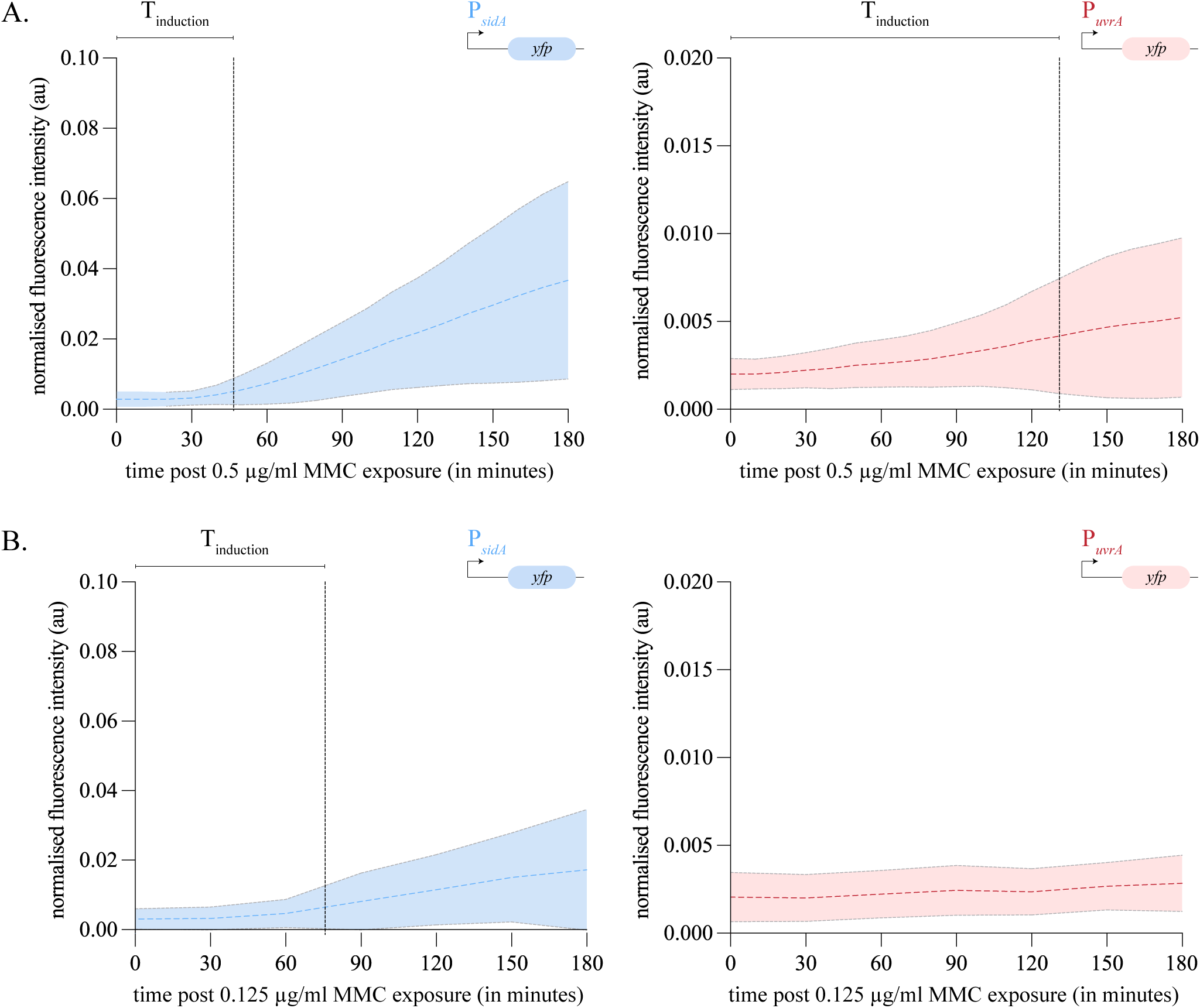
The *Caulobacter* SOS response exhibits temporal hierarchy. **(A)** Fluorescence intensity normalized to cell area for P*_sidA_yfp* [top] and P*_uvrA_yfp* [bottom] cells over 3 hours of MMC exposure. Images were taken every 10 min and intensity over time traces are shown. Dashed line indicates the mean and the shaded region indicates the standard deviation. Time to induction (T_ind_) is indicated on the graph. (0.5 μg/mL, n=25). **(B)** As (A) for cells treated with 0.125 μg/mL MMC, except for an imaging interval of 30 min (n=20). At these low doses induction of *yfp* from the *uvrA* promoter is not detected.

**Figure S3.**
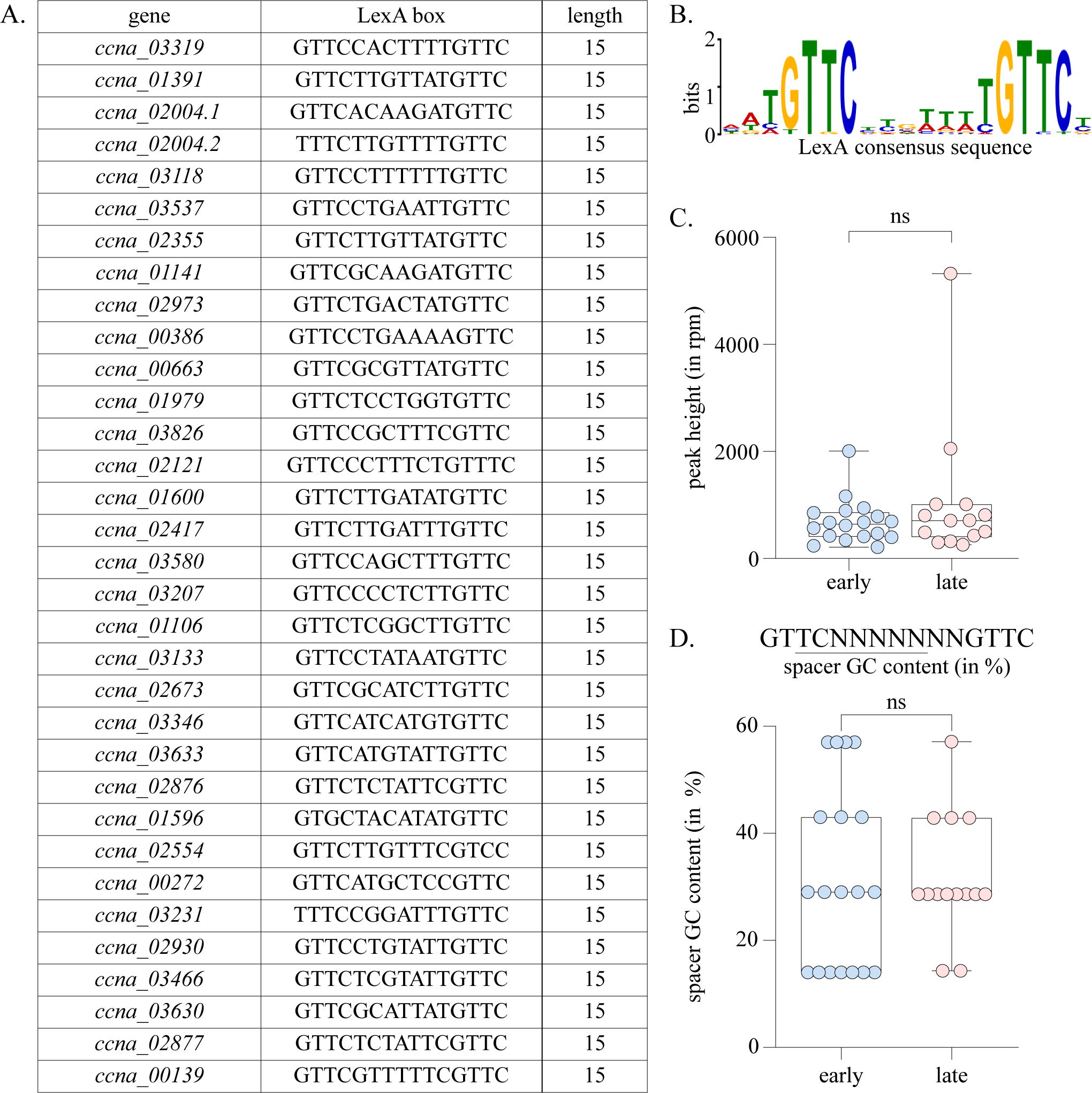
LexA properties are not predictive of the temporal hierarchy in SOS response induction. **(A)** List of LexA box sequence and their box length (in bp) for promoters of all *Caulobacter* SOS response genes analysed in present study. **(B)** Binding consensus motif for LexA, based upon the promoters of the *Caulobacter* SOS response genes. **(C)** Box and scatter plot indicating peak height derived from LexA ChIP-seq data for early (blue) and late (red) SOS response genes. Mann Whitney test, n.s. – not significant. **(D)** Box and scatter plot indicating %GC content of the LexA box spacer for the CDS of early (blue) and late (red) SOS response genes. Mann Whitney test, n.s. – not significant.

**Figure S4.**
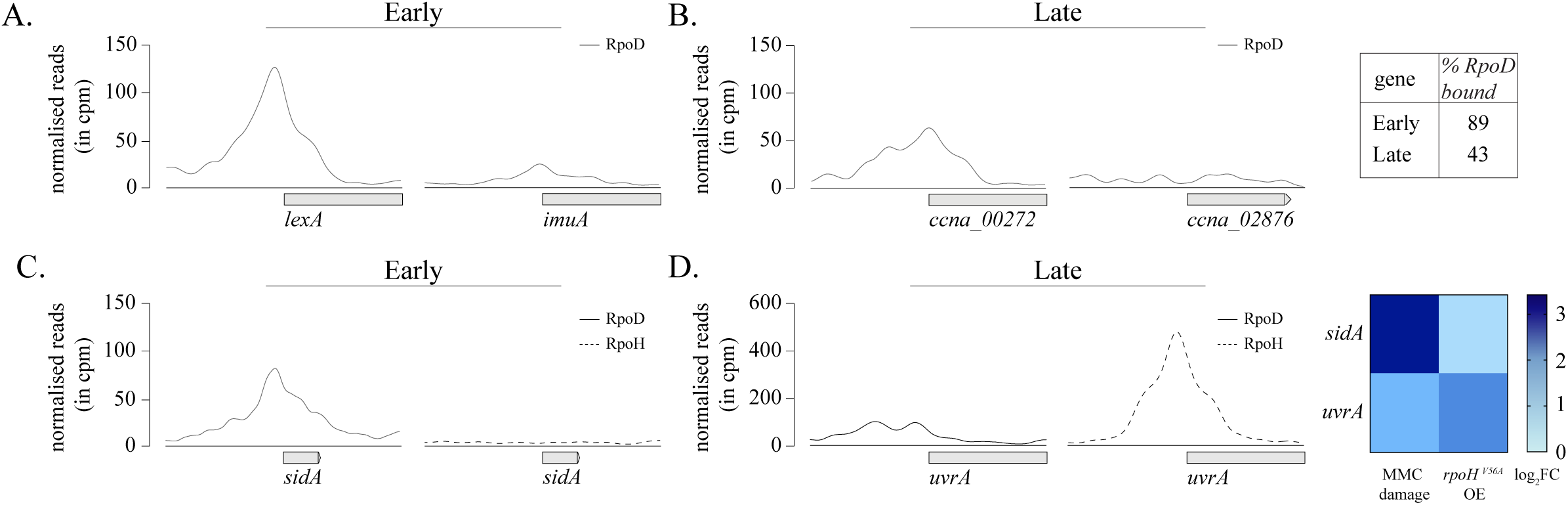
Intrinsic promoter properties contribute to temporal hierarchy in SOS response gene induction. **(A-B)** ChIP-seq profiles for RpoD enrichment 500bp upstream and downstream of the CDS for early [left] and late [right] SOS response genes. % SOS response genes bound by RpoD are indicated adjacent to the ChIP profiles. ChIP-seq data were obtained from the GEO database (GSE73925) [35]. **(C-D)** ChIP-seq profiles for RpoD and RpoH enrichment 500bp upstream and downstream of the CDS of early (*sidA*) and late (*uvrA*) SOS response genes. ChIP-seq data were obtained from GEO database (GSE73925) [35]. Heat map of log_2_FC values from RNA-seq experiments for *Caulobacter* SOS response genes upon *rpoH^V65A^* over-expression is indicated adjacent to the ChIP profiles. RNA-seq data were obtained from GEO database (GSE102372) [36]. For comparison log_2_FC values for the same genes under 40 min of MMC damage (0.25 μg/mL) are shown.

## Supplementary tables

Table S1: Strains used in present study

Table S2: Plasmids used in present study

Table S3: Oligos used in present study

Table S4: Genes induced at 20 and 40 min post MMC damage. Gene expression profile for these genes is provide for *recA* deletion and *lexAsidA* deletion cells as well

Table S5: List of *Caulobacter* SOS response genes and their gene annotations

Table S6: σ^70^ or σ^32^ binding at SOS promoters categorized as per their temporal properties

Table S7: Summary of coefficient of variation (CV) values for P*_sidA_yfp* and P*_uvrA_yfp* promoter fusions in wild type (+/- MMC damage) and *ΔlexA* background.

